# Characteristics of *N*^6^-methyladenosine modification during sexual reproduction of *Chlamydomonas reinhardtii*

**DOI:** 10.1101/2022.03.26.485907

**Authors:** Ying Lv, Fei Han, Mengxia Liu, Ting Zhang, Guanshen Cui, Jiaojiao Wang, Ying Yang, Yun-Gui Yang, Wenqiang Yang

## Abstract

The unicellular green alga *Chlamydomonas reinhardtii* (referred to as *Chlamydomonas* throughout) is an ideal model organism which possesses both plant and animal attribute for studying the fundamental processes such as photosynthesis, sexual reproduction and life cycle *etc*. *N*^6^-methyladenosine (m^6^A) is the most prevalent mRNA modification and plays important roles during sexual reproduction in animals and plants. However, the pattern and function of m^6^A modification during *Chlamydomonas* sexual reproduction is still unknown. Here, we performed transcriptome and MeRIP sequencing on the six samples from different stages during sexual reproduction of *Chlamydomonas* life cycle. The results showed that m^6^A occurs widely at the main motif of UGKAM (K= U/G, M= A/C) in *Chlamydomonas* mRNA. Moreover, m^6^A peak in *Chlamydomonas* mRNA is mainly enriched in the 3’UTR region and negatively correlated with the abundance of the transcripts at each stage. Especially, genes in microtubule-associated pathway showed a significant negative correlation between gene expression level and m^6^A level, indicating the influences of m^6^A modification on sexual reproduction and life cycle of *Chlamydomonas* through regulating microtubule-based movement. In summary, our findings first demonstrate the distributions and the functions of m^6^A modification in *Chlamydomonas* and provide new insights into the understandings of m^6^A modification in the process of sexual reproduction in other plant organisms evolutionarily.

## Introduction

In eukaryotes, sexual reproduction consists of two sequential events: haploid gametes fuse to form diploid zygotes after fertilization, and diploid zygotes produce haploid progenies by meiosis [1, 2]. Unlike animals and plants, the unicellular green alga *Chlamydomonas* has unicellular haploid body and diploid body like the gametophyte and sporophyte of land plants, respectively [1, 3–5]. *Chlamydomonas* is known as “Green Yeast”, and has been introduced as a model organism to study fundamental processes such as photosynthesis, nutrient metabolism, flagella biology, cell cycle and sexual reproduction [5–11]. Because *Chlamydomonas* possesses both animal and plant feature, the study of cell cycle and sexual reproduction in *Chlamydomonas* is important in evolution [1]. Many studies have shown that mitotic cell cycle has a long G1 phase and rapidly alternating S/M phases which facilitates *Chlamydomonas* to produce 2^n^ (n=1∼5) daughter cells in one cell cycle [9, 12]. In *Chlamydomonas*, the cell cycle and sexual reproduction is mainly controlled by two major cyclin dependent kinases of CDKA1 and CDKB1 [13, 14], and other critical proteins reported previously [15, 16]. The *Chlamydomonas* vegetative cells can be induced to produce two kinds of isogametes: mating type minus (mt+) and mating type plus (mt-) by nitrogen deficiency [10, 17–19], and many mating type specific genes are induced during this process [20–22]. Gamete-specific agglutinins encoded by the mt+ specific gene (*SAG1*) and the mt-specific gene (*SAD1*) facilitate the interactions between two different mating type gametes [23]. Different mating type gametes adhere together to initiate zygote formation procedure including the increase cAMP levels as signal, flagella tip activation, loss of cell wall and mating apparatus activation accompanied with actin polymerization [24–29]. After clumping together, the mating type specific structures are formed and this process is controlled by FUS1 (mt+) and GCS1/HAP2 (mt-), respectively [28, 30]. Simultaneously, the expression of zygote-specific genes such as *EZYs* is up-regulated [15], and the transcription factors GSP1 (mt+) and GSM1 (mt-) accumulate to regulate the transition from haploid cells to the diploid zygotes [3, 4, 10, 22, 31].

In eukaryotes, RNA modifications are very important for the fate determination of RNAs [32–34]. As the most prevalent regulator on eukaryotic mRNAs, m^6^A modification functions in mRNA alternative splicing, nuclear export, stability, translation and degradation [35–43]. In mammals, m^6^A modification is catalyzed by a large RNA methyltransferase complex (MTase) as writer protein, which is composed of methytransferase-like 3 (METTL3), methyltransferase-like 14 (METTL14), Willms’ tumor 1-associating protein (WTAP), Virilizer like m^6^A methyltransferase associated protein (VIRMA), Cbl photo oncogene like 1 (CBLL1), RNA-binding protein 15/15B (RBM15/15B) and Zinc finger CCCH domain-containing protein 13 (ZC3H13) [44–52]. The removal of methyl groups is performed by two RNA demethylases as eraser proteins, Fat mass and Obesity-associated protein (FTO) and Alkylated DNA repair protein alkB homolog 5 (ALKBH5) [53–60]. Furthermore, the function of m^6^A modification in mammalian mRNA metabolic processes mainly depends on the diverse reader proteins, including YTH domain-containing family (YTHDF1-3, YTHDC1-2), IGF2 mRNA binding protein family (IGF2BP1-3), and heterogeneous nuclear ribonucleoproteins (HNRNPC, HNRNPG, HNRNPA2B1) [36, 37, 39, 40, 42, 43, 61–68]. Malfunction of these proteins causes disorder in m^6^A modification and further affects spermatogenesis and embryonic development in animals [60, 69–71]. Consistently, m^6^A modification is also detected in plants, and the animal counterpart of writers, erasers and readers have also been identified to play critical roles [72–83]. Functional studies showed that m^6^A is also important for embryo development and sporogenesis in plants [6, 75, 77, 84–87]. These findings suggest that m^6^A modification may have conserved critical roles in sexual reproduction in mammals and plants [60, 69–71, 75, 77], whether and how m^6^A modification is involved in sexual reproduction and life cycle regulation of *Chlamydomonas* remains unknown.

Here, we first performed MeRIP sequencing to depict the m^6^A modification landscapes on six samples during sexual life cycle of *Chlamydomonas* with two biological replicates (mt+ vegetative cells, mt-vegetative cells, mt+ gametes, mt-gametes, zygotes on 1 day and 7 days), and RNA-seq was conducted simultaneously to analyze the associated transcriptional variation. The results showed that m^6^A peaks appear widely in *Chlamydomonas* mRNA, while m^6^A peaks are mainly enriched in 3’ UTR region of mRNA. UGKAM (where K is U/G, A is m^6^A, M is A/C) is the main motif of m^6^A modification peak with slight differences among different stages.

Moreover, the combined analyses of MeRIP-seq and RNA-seq showed that *Chlamydomonas* m^6^A modification is negatively correlated with the abundance of the transcripts, especially the genes in microtubule-associated pathways showed significantly opposite performance between gene expression and m^6^A modification level, suggesting that m^6^A modification exerts functions through regulating microtubule-based movement during sexual reproduction in *Chlamydomonas* life cycle. Finally, CrMETTL3 and CrMETTL14 may function as potential m^6^A methyltransferases in *Chlamydomonas* responsible for m^6^A formation. Overall, the findings reveal the distribution features of m^6^A modification and its potential regulatory functions during sexual reproduction and life cycle in *Chlamydomonas*.

## Results

### Features of critical periods during life cycle of *Chlamydomonas*

To examine the dynamics of m^6^A modification during sexual reproduction of *Chlamydomonas* life cycle, six samples of key periods [mt+ vegetative cells, mt-vegetative cells, mt+ gametes, mt-gametes, zygotes on 1 day (1d zygotes) and zygotes on 7 days (7d zygotes)] were collected for MeRIP-seq analyses (**Figure 1A**). Asexual reproduction mainly undergoes vegetative growth and mitosis, and the vegetative cells at this stage have classical morphological characteristics: around 5∼7 μm in diameter, with two flagella, one cup-shaped chloroplast, an eyespot, a nucleus and other organelles (**Figure 1B**). The vegetative cells are induced to form gametes by nitrogen starvation or blue light, which is called gametogenesis [88]. The gametogenesis of *Chlamydomonas* is presumed to be a process of stress response, as the gametes show high motility and low activity of photosynthesis. It should be noted that the size of gametes is much smaller than that of vegetative cells (**Figure 1C**). When the induced gametes of the opposite mating type are mixed together to form zygotes, the mating responses are triggered rapidly following with the adhesion of mating type specific agglutinins on the surface of flagella [16]. Compared with the vegetative cells, the size of zygote without flagella is larger with the thicker cell wall (**Figure 1D and 1E**). After 1 day in the light and 6 days in the dark, the zygote develops into zygospore which is more resistant to the various of stresses. With favorable environments, the zygospores germinate, and meiosis occurs to release haploid vegetative cells.

**Figure 1.**
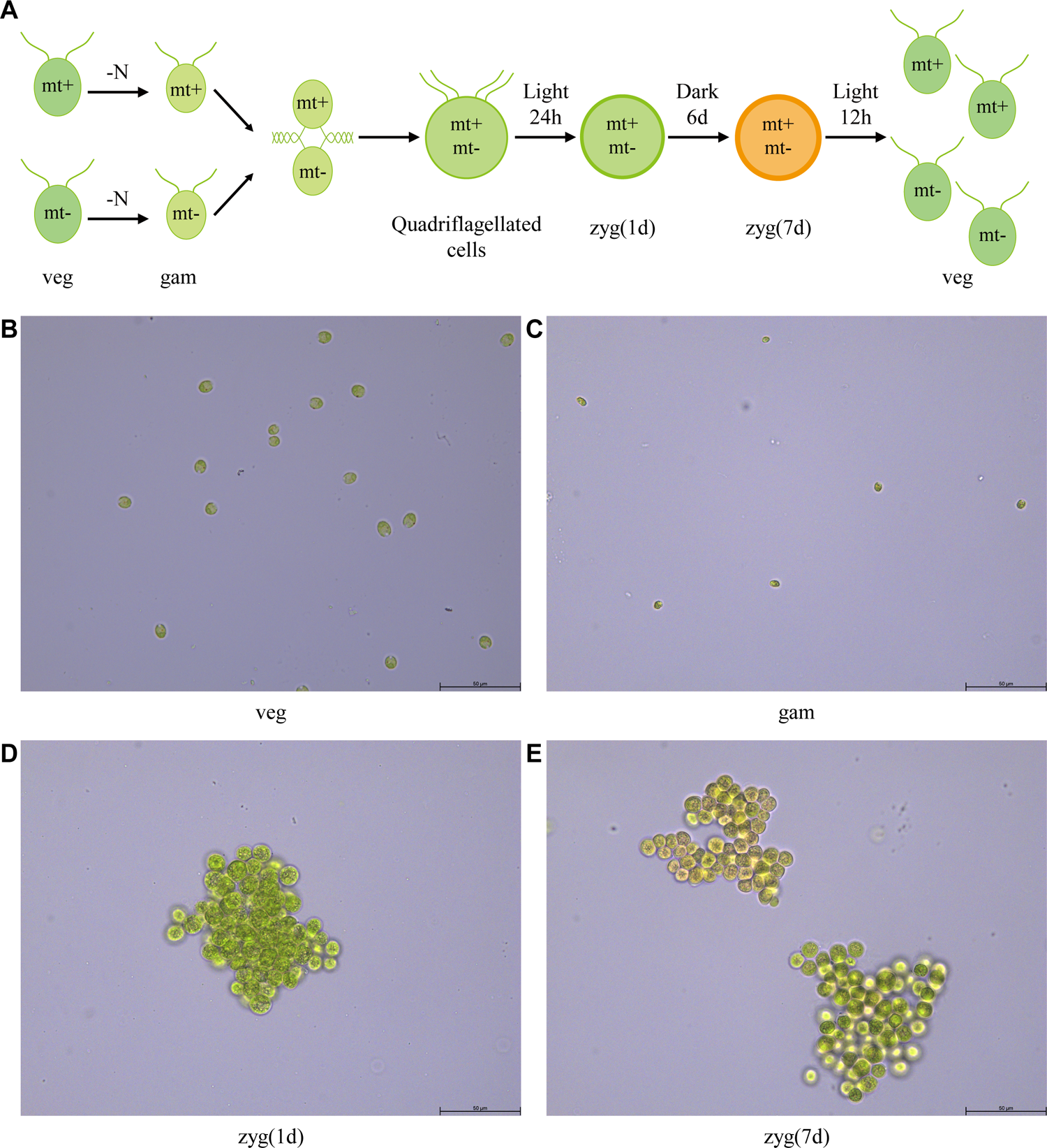
Sexual life cycle of *Chlamydomonas* and the features of critical periods. **A.** Overview of the life cycle of *Chlamydomonas.* Vegetative cells can differentiate into gametes in response to nitrogen deficiency; zygotes can be formed from the mating of gametes with different mating types under light and eventually be mature in a few days without light; and mature zygotes conversely are able to germinate to form progenies of 4 daughter cells through tetrad by the addition of nitrogen and light. **B-E.** Image to show the life cycle at different stages. vegetative cells (B), gametes (C), 1d zygotes (D), and 7d zygotes (E) under 400× light microscope, respectively.

### m^6^A modification shows dynamic changes during the process of sexual reproduction of *Chlamydomonas*

RNA m^6^A modification has been shown with critical roles in sexual reproduction in mammals and plants [60, 69–71, 75, 77]. To examine the levels of m^6^A modification during sexual reproduction of *Chlamydomonas* life cycle, Dot-blot was performed at different stages during the sexual reproduction as described above, which showed that the level of m^6^A modification is down-regulated in gametogenesis and up-regulated in zygote development (**Figure 2A**). Based on the presence of predicted RNA adenosine methylase domains (MTA70) and full-length human METTL3 and METTL14 protein sequences, four candidates of *Chlamydomonas* m^6^A methyltransferases were found from *Chlamydomonas* Phytozome (https://phytozome-next.jgi.doe.gov/), which are CrMETTL3 (Cre06.g295600), CrMETTL14 (Cre01.g050600), CrMT1 (Cre06.g288100), and CrMT2 (Cre10.g452300) (**Figure 2B**). Among them, CrMT1 and CrMT2 show lower homology to human METTL3 and METTL14. Quantitative real-time PCR showed that *CrMETTL3* and *CrMETTL14* expression levels consistently declined in gametes with different mating types and increased in 1d zygotes then declined again in 7d zygotes (**Figure 2C**), suggesting that m^6^A modification may participate in regulating the sexual reproduction and change dynamically in *Chlamydomonas*.

**Figure 2.**
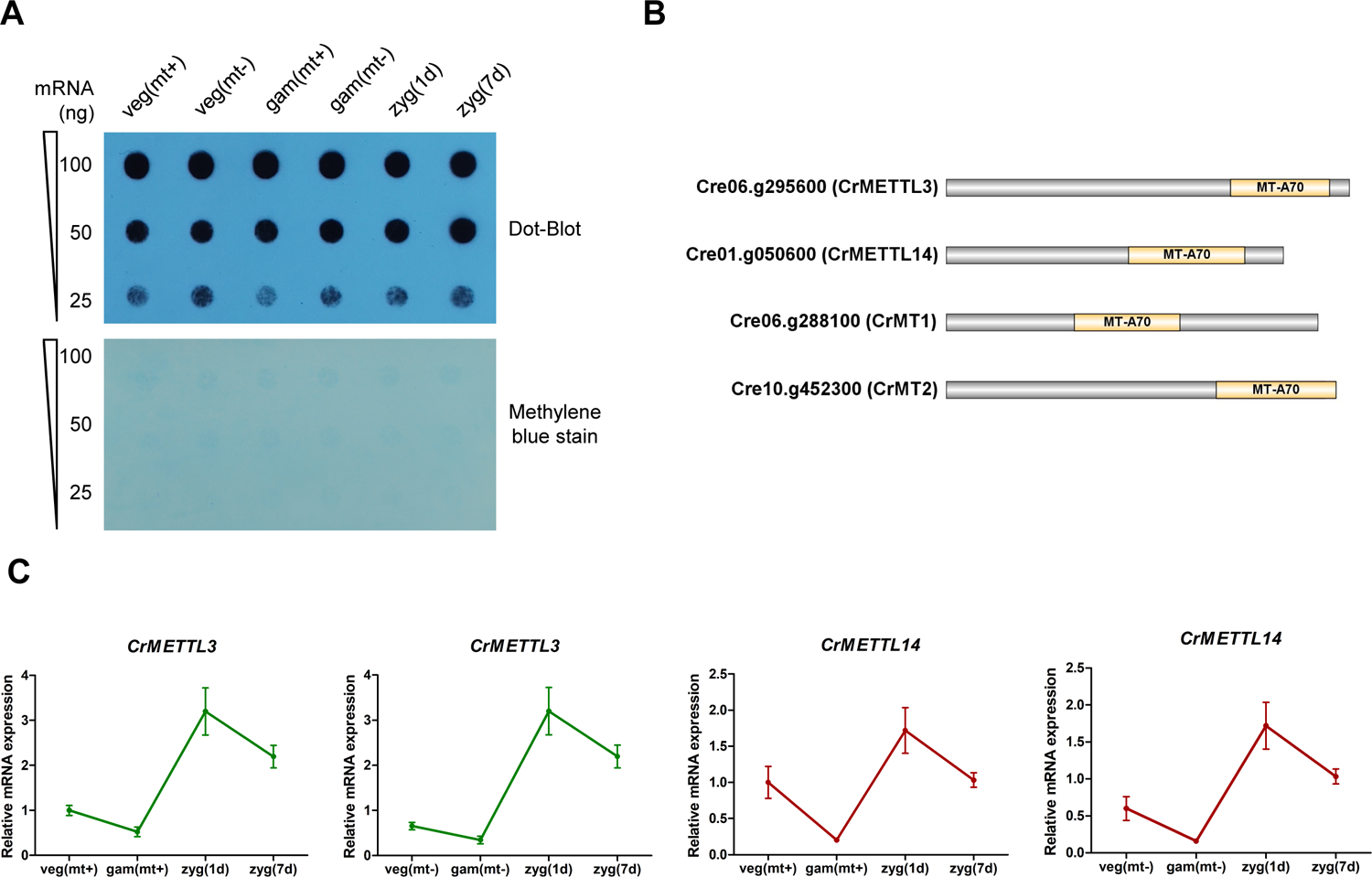
mRNA m^6^A methylation shows dynamic changes during the life cycle of *Chlamydomonas*. **A.** The overall levels of m^6^A mRNA modification detected by dot-blot assays using a specific anti-m^6^A antibody (upper panel), and the methylene blue staining to show loading control (lower panel). **B.** Schematic representations of the putative candidates of m^6^A methyltransferases in *Chlamydomonas*. MT-A70 represents the conserved motif. **C.** qRT-PCR to show the relative abundance of transcripts of the putative genes encoding m^6^A methyltransferases in *Chlamydomonas*. The internal control was *CBLP* gene. Error bars represent standard deviation of three biological replicates. Veg (mt+): mating type plus vegetative cells; veg (mt-): mating type minus vegetative cells; gam (mt+): mating type plus gametes; gam (mt-): mating type minus gametes; zyg (1d): 1d zygotes; zyg (7d): 7d zygotes.

### Overview of m^6^A methylome in *Chlamydomonas*

To explore the potential role of m^6^A modification in regulating *Chlamydomonas* sexual reproduction, MeRIP-seq was performed on the samples from different stages to compared their transcriptome-wide m^6^A methylome. We detected m^6^A peaks with high confidence (FDR<0.05) using exomePeak. 24416, 25656, 23288, 23836, 24403 and 25057 m^6^A peaks within 13255, 13367, 12969, 13313, 13564 and 13720 transcripts were identified from different stages, respectively (**Figure 3A** and **Table S1**). The distribution pattern of m^6^A modification along the transcripts were analyzed, and the results of metagene profiles revealed that m^6^A deposition was primarily enriched in 3’UTR (**Figure 3B**), which is interestingly consistent with the m^6^A distribution patterns in rice, potato and maize [85, 86, 89]. We then analyzed the distribution of m^6^A peaks within four non-overlapping regions: intergenic region, 5’ UTR, coding sequence (CDS), and 3’UTR. Among them, m^6^A peaks appeared to be greatly enriched in 3’UTR segment (**Figure 3C**), with 62 ∼67% of the peaks from different samples fell into this region. Furthermore, the distribution density plot of m^6^A peaks across the exon length exhibited that m^6^A peaks tend to occur within the exons around 760 bp in length (**Figure 3D**), indicating that the m^6^A modification ends to be catalyzed on the long exons while the average length of exons in *Chlamydomonas* is 376.62 nucleotides.

**Figure 3.**
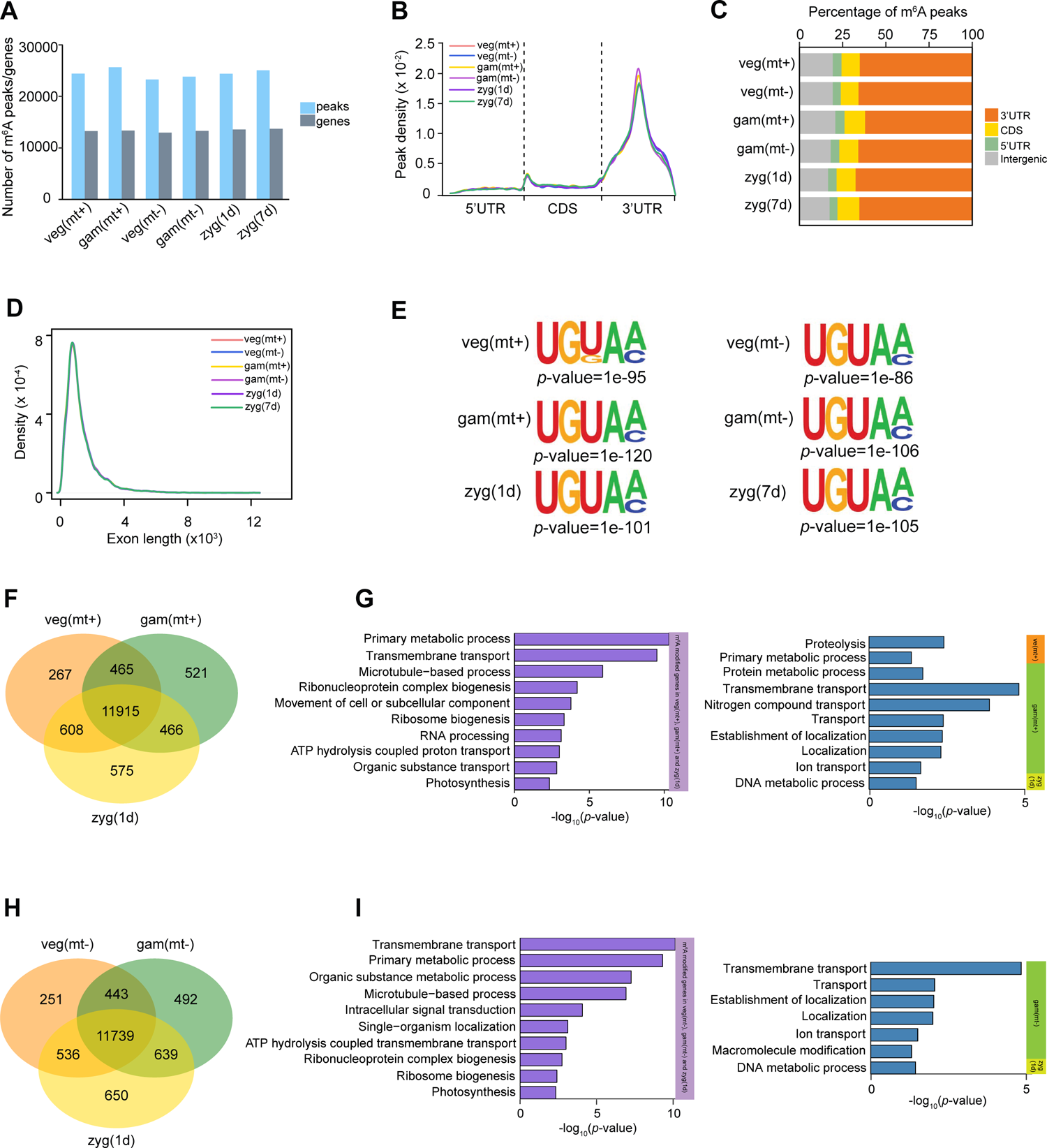
Overview of m^6^A methylome in *Chlamydomonas*. **A.** Histograms to show numbers of the detected m^6^A peaks and methylated genes at each stage of the *Chlamydomonas* life cycle. **B.** Box charts to show the methylation level of m^6^A peaks in the various of tested samples. **C.** Metagene profiles to show the distribution of m^6^A peaks along the transcripts composed of three rescaled non-overlapping segments (5′UTR, CDS, and 3′UTR). UTR, untranslated region; CDS, coding sequence. **D.** Stacked bar charts showing the percentage of m^6^A peaks within distinct RNA sequence types. **E.** Density plots showing the distribution of m^6^A peaks across the length of exon. **F.** Top sequence motifs identified within m^6^A peaks. **G, I.** Venn diagram showing the overlap of m^6^A modified genes among vegetative cells, gametes and 1d zygotes with mating type plus. **H, J.** Bar plot exhibiting Gene Ontology (GO) enrichment of m^6^A commonly modified genes (left) and specifically modified genes (right) among vegetative cells, gametes and 1d zygotes with mating type minus.

To identify the consensus sequence and enrichment of m^6^A peaks appeared in the transcriptome, motif search was performed for high confident m^6^A peaks using HOMER. The RRACH (R is A/G, A is m^6^A, and H is A/C/U) motif is commonly observed in mammals [90, 91], while it was not significantly found in the m^6^A peaks of this study. The UGKAM (K is U/G, A is m^6^A, and M is A/C) motif (**Figure 3E**) was significantly identified and top ranked in all of the detected stages including vegetative cells, gametes, and 1 day and 7 days zygotes. This motif is in general similar to the motifs found in rice, tomato and maize seedlings [85–87].

We next compared the transcriptome-wide m^6^A methylome during sexual reproduction. 11915 m^6^A modified genes were shared among mt+ vegetative cells, mt+ gametes and 1d zygotes, and 11739 m^6^A modified genes were shared among mt-vegetative cells, mt-gametes and 1d zygotes, respectively (**Figure 3F and 3H**). Only less than 10% of the specific m^6^A modified genes were detected in all stages. To further analyze the regulatory mechanisms of m^6^A modification during sexual reproduction, we then performed Gene Ontology (GO) analysis of the genes with m^6^A modification. The common m^6^A modified genes in mt+ vegetative cells, mt+ gametes and 1d zygotes were enriched in primary metabolic process, transmembrane transport and RNA processing (**Figure 3G**), implying that m^6^A is essential for the basic life activities in *Chlamydomonas*. m^6^A modification was also found to be related to microtubule-based process and photosynthesis, which influence the gametogenesis and zygotes development during the sexual reproduction. Interestingly, the enriched processes by common m^6^A modified genes in mt-vegetative cells, mt-gametes and 1d zygotes were similar to the previous findings in mating type plus (**Figure 3I**). Genes with specific m^6^A methylation at different stages were related to proteolysis and DNA metabolic process (**Figure 3G and 3I**), indicating that m^6^A methylation generally appears in various stages and is involved in the regulation of important processes during the sexual reproduction of *Chlamydomonas*.

### m^6^A modification is generally negatively correlated with gene expression level

m^6^A has been proven to regulate the stability of mRNA [40, 76, 78]. According to the m^6^A methylome at different stages, we further examined the gene expression level to investigate the m^6^A regulation in mRNA abundance during the sexual reproduction. We determined the mRNA abundance as previously described and obtained transcriptome-wide RNA expression map with strong correlation between biological replicates (**Figure 4A and S2A**). Briefly, 12465, 11043, 13229, 11166, 13241 and 13347 stable expressed transcripts were obtained in the various of tested samples, respectively (**Figure 4B** and **Table S2**). The expression of some stage-specific genes was also determined (**Figure S2B**), including the well-known *GSP1* (Cre02.g109650) and *GSM1* (Cre08.g375400), which encode transcription factors specifically expressed in mating type plus and mating type minus gametes, respectively, and are involved in the development of zygotes [20–22]. Moreover, the early zygote-specific genes were also identified, including *EZY9* (Cre06.g304500), *ZYS2/MAW1* (Cre07.g325812) and *EZY3* (Cre11.g482650) in 1d zygotes [15].

**Figure 4.**
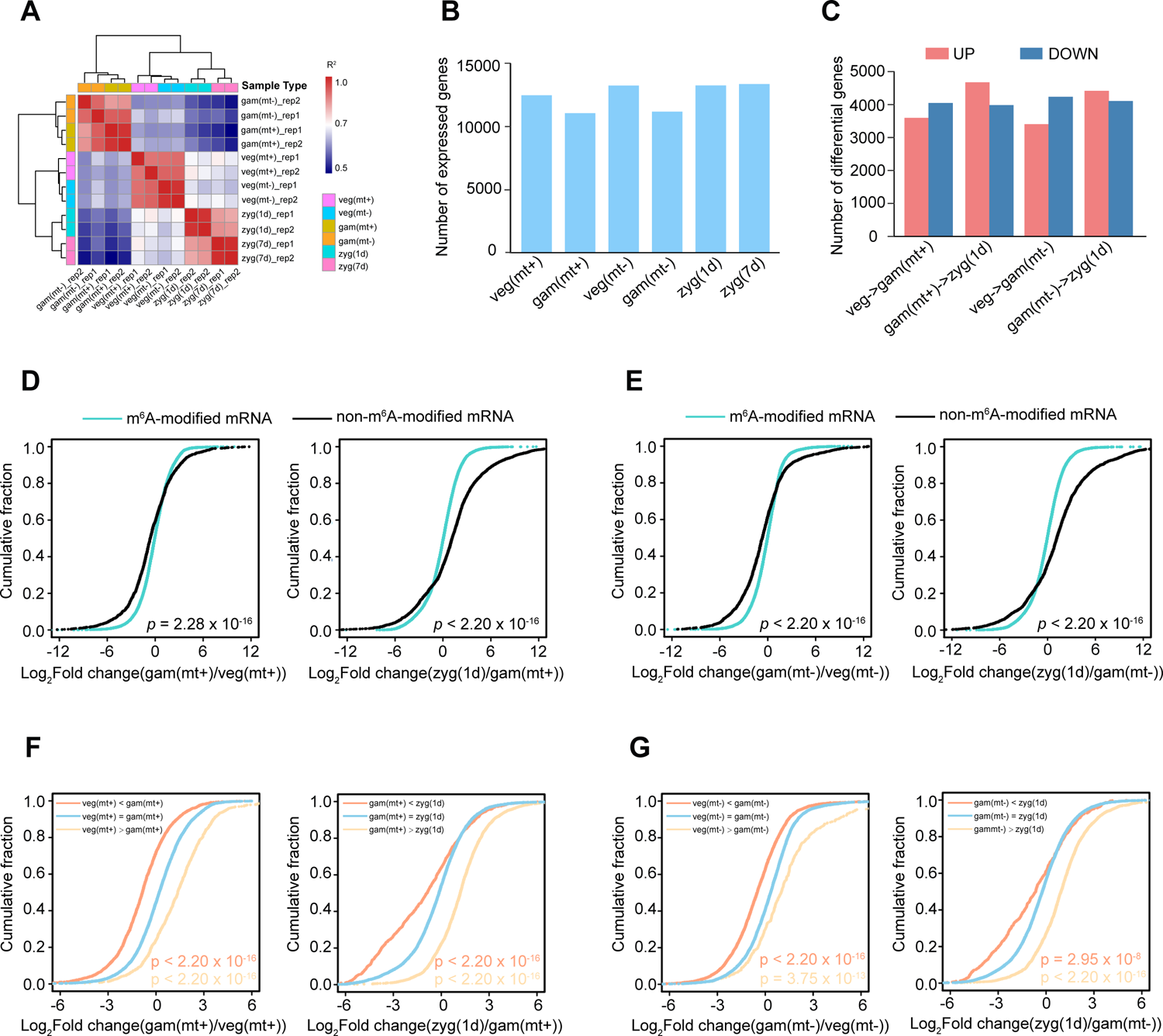
m^6^A modification is generally negatively correlated with gene expression level. **A.** Heatmap showing the high correlation among replicates based on the expression matrix. **B.** Histograms to show the number of genes expressed at each stage of the *Chlamydomonas* life cycle. **C.** The number of differentially expressed (up-regulated and down-regulated) genes at different stages. **D-E.** Cumulative distribution displaying the expression level changes in mRNAs with or without m^6^A methylation in mating type plus (**D**) vegetative cells, gametes and zygotes and mating type minus (**E**) vegetative cells, gametes and zygotes. **F-G.** Cumulative distribution displaying the expression level changes in mRNA classified by m^6^A methylation level in mating type plus (**F**) vegetative cells, gametes and zygotes and mating type minus (**G**) vegetative cells, gametes and zygotes.

We next analyzed the differentially expressed genes to investigate the transcriptional changes during the sexual reproduction. A total of 7645 and 7640 transcripts were differentially expressed (|log_2_(fold change)| > 1; FDR < 0.01) between vegetative cells and gametes with mating type plus and minus (**Figure 4C, Table S3**), among which 4048 and 4235 genes were down-regulated in mating type plus and minus gametes, respectively. Additionally, we identified 8525 differentially expressed genes between mating type plus gametes and zygotes, and 8701 genes in mating type minus gametes and zygotes. Among them, 4675 and 4418 genes exhibited higher expression level in zygotes than that in mating type plus and minus gametes, respectively. We next compared the differentially expressed genes between mating type plus and minus. The results (**Figure S2C**) showed that more than 60% differentially expressed genes were shared between different mating types during the transitions from vegetative cells to gametes and from gametes to zygotes.

To study whether the changes of mRNA abundance were influenced by m^6^A modification during the sexual reproduction, the association of m^6^A with mRNA abundance were analyzed. Cumulative distribution showed a significant difference between the mRNA abundance of m^6^A modified genes and non-m^6^A modified genes during the mating type plus gametogenesis and mating type plus gametes to 1d zygotes (**Figure 4D**). Similar tendency was also found in the processes 1d of mating type minus gametogenesis and mating type minus gametes to 1d zygotes (**Figure 4E**), indicating that m^6^A methylation is likely affect mRNA abundance during sexual reproduction in *Chlamydomonas*.

To further elucidate the relationship between the changes in mRNA abundance and m^6^A methylation levels, the m^6^A modified genes were classified into three types: genes with higher methylation levels, lower methylation levels and stable methylation levels. In the process from mating type plus vegetative cells to gametes, the changes of abundance in the three types of m^6^A modified genes showed great contrast variability. Compared with genes with stable methylation, the genes with higher m^6^A methylation levels in gametes tend to be down-regulated, while demethylated genes generally exhibit significantly higher abundance (**Figure 4F**). These results suggested that the mRNA abundance is negatively regulated by m^6^A methylation in the plus gametogenesis. The negative correlation between methylation level and mRNA abundance was also observed in other samples from different stages, including minus gametogenesis (**Figure 4G**), plus gametes to 1d zygotes (**Figure 4F**) and minus gametes to 1d zygotes (**Figure 4G**), suggesting that m^6^A modification negatively regulate mRNA abundance during the sexual reproduction of *Chlamydomonas*.

### m^6^A is involved in the sexual reproduction through regulating microtubule-based movement

To further explore how m^6^A plays a role in the sexual reproduction, we investigated the biological process related to sexual reproduction and analyzed whether m^6^A could negatively regulate the expression levels of key genes involved. Gene Ontology (GO) analyses showed most of the down-regulated genes during the mating type plus and minus gametogenesis were enriched in photosynthesis-associated processes (**Figure S3A and S3C**). We speculate that the vegetative cells need to reduce the photosynthesis in response to nitrogen starvation and undergo gametogenesis. Most of the up-regulated genes from mating type plus vegetative cells to gametes were specifically enriched in microtubule-based process and cilium organization (**Figure 5A**). Previous researches have reported that genes associated with microtubule-based process encodes kinesin and dynein proteins, which mediated intraflagellar transport system allowing agglutinins to be transported to flagellar membrane surfaces and also were involved in flagella assembly to regulate gametogenesis [16, 92–94]. Proteome analysis of male and female gametocytes of plasmodium reveals that kinesin and dynein are proteins expressed in a sex-specific manner [95]. Among the specifically up-regulated genes related to microtubule-based process in plus gametes (**Figure 5B**), *KIN9-3* (Cre10.g427750) encodes a kinesin protein, which showed significantly lower m^6^A methylation levels, indicating that *KIN9-3* is involved in flagellar assembly and associated with sexual specific flagellum formation during the plus gametogenesis [96] (**Figure 5C**). In addition, qRT-PCR was applied to confirm the upregulation of *KIN9-3*, *DHC1* (Cre12.g484250), and *DHC8* (Cre16.g685450) (**Figure 5C**).

**Figure 5.**
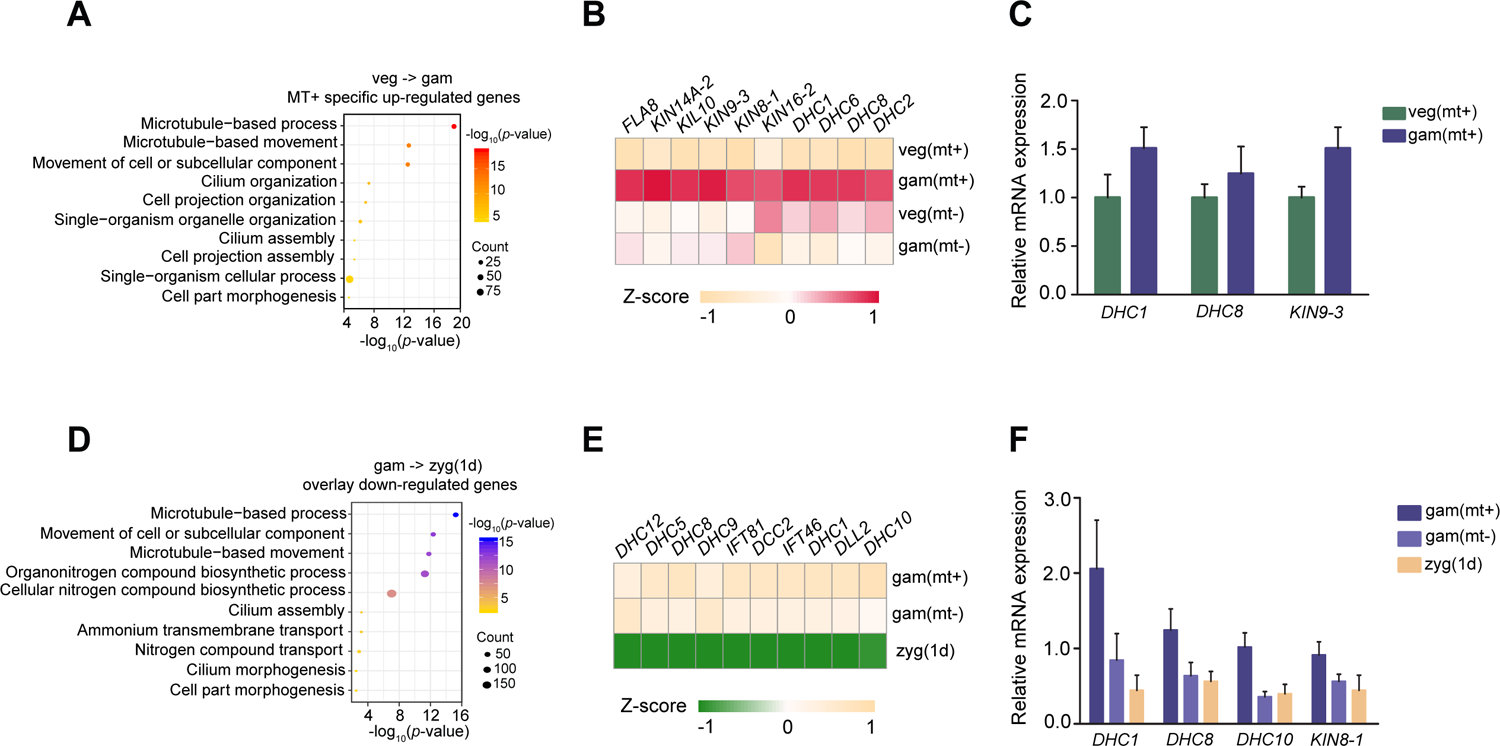
m^6^A is involved in the life cycle through regulating microtubule-based movement. **A.** Gene Ontology (GO) enrichment of the up-regulated genes in gam (mt+) during the gametogenesis. **B.** Heatmap of up-regulated genes associated with microtubule-based process in gametogenesis. **C.** qRT-PCR to validate the target genes regulated by m^6^A in gametogenesis. The internal control was *CBLP* gene. Error bars represent standard deviation of three biological replicates. **D.** GO enrichment of the down-regulated genes in 1d zygotes during the process of gametes fusion. **E.** Heatmap of down-regulated genes associated with microtubule-based process in the process of gametes fusion. **F.** qRT-PCR to validate the target genes regulated by m^6^A during gametes fusion. The internal control was *CBLP* gene. Error bars represent standard deviation of three biological replicates.

After gametes fusion, light exposure over a period of time ensures zygotes formation and maturation. We detected the global increase in the expression level of photosynthesis-associated genes in the process from the plus and minus gametes to zygotes, indicating the diploid zygotes may gain energy through photosynthesis (**Figure S3B and S3C**). More interestingly, we found that genes with lower expression in zygotes in both plus and minus gametes participate in the microtubule-based process, movement of cell or cellular component and cilium organization, indicating that the motility was reduced in mature zygotes (**Figure 5D and 5E**). The identified down-regulated microtubule-related genes with higher m^6^A modification levels are considered as candidate genes regulated by m^6^A during the zygotes formation. Among them, *DHC1*, *DHC8, DHC10* (Cre14.g624950) and *KIN8-1* (Cre13.g602400), as the key important genes in microtubule-based process, were further verified (**Figure 5F**) [97]. These results suggest that KINs and DHCs are critical for the fulfillment of the sexual reproduction in *Chlamydomonas*.

## Discussion

As the most prevalent mRNA modification in eukaryotes [98], m^6^A modification is involved in many essential biological processes, including cell fate determination [69, 99] and embryonic development [69, 100, 101]. In addition, m^6^A modification functions during embryo development and sporogenesis in plants [6, 75, 77, 84]. These findings suggest that m^6^A modulation of mRNA expression may play important roles in sexual reproduction of both animals and plants. *Chlamydomonas* is an excellent model organism for studying sexual reproduction and life cycle, because it has both characteristics of plants and animals [6]. However, the role and function of m^6^A during this process in *Chlamydomonas* remained unclear.

During sexual reproduction and life cycle of *Chlamydomonas*, the cells undergo the transition of haploid phase to diploid phase. The vegetative cells and gametes are haploid while zygotes are diploid. In this study, we performed MeRIP-seq on six samples with duplicates from different stages during *Chlamydomonas* sexual reproduction and life cycle and compared their transcriptome-wide m^6^A methylome.

Our analyses reveal that m^6^A peaks is remarkedly enriched in 3’UTR and the distribution pattern of m^6^A is similar among all of the samples. Interestingly, the most enriched sequence motif identified within m^6^A peaks is UGKAM but not the canonical RNA motif of RRACH [42, 43], and the UGKAM motif is found similar to the motifs characterized in rice, tomato and maize seedlings. Moreover, by combining MeRIP-seq and RNA-seq, we discovered that m^6^A level is generally negatively correlated with the abundance of the corresponding mRNAs. Collectively, our data reveal the dynamic changes of m^6^A methylation and the negative correlation between methylation levels with stage-specific transcripts during sexual reproduction and life cycle of *Chlamydomonas*, suggesting a regulatory role of m^6^A in sexual reproduction. Previous studies have reported that kinesin and dynein proteins are associated with microtubule-based process and are important sex-specific expressed proteins, which mediate intraflagellar transport system and also are involved in flagellar assembly to regulate gametogenesis [16, 88, 89]. In our study, microtubule-based process and cilium organization from plus vegetative cells to gametes are also specifically accumulated. Among the specifically up-regulated genes in mating type plus gametes, *KIN9-3*, *DHC6* and *KIN16-2* show significantly lower m^6^A methylation levels and closely related to microtubule-based process during the plus gametogenesis.

Additionally, the genes of *DHC1*, *DHC8, DHC10* and *KIN8-1,* with lower expression and higher m^6^A methylation levels in zygotes are also shown to be involved in the microtubule-based process. All of these data suggest that m^6^A modification potentially participates in sexual reproduction and life cycle through regulating the abundance of transcripts involved in microtubule-based movement. It is noteworthy that the expression of photosynthetic-related genes including PSI, PSII assembly and those involved in light harvesting and oxidation-reduction cycle decreases during gametogenesis (**Figure S3C**), which is possibly related to the nitrogen deficiency treatment to activate gametogenesis program. Similar results have also been observed in the previous study witch showed the similar trend of GreenCut2 [15, 102]. Limitation of nitrogen leads decreased activity of photosynthesis, down-regulation of carbon assimilation and chlorophyll biosynthesis in *Chlamydomonas* [103]. The limited substrate and energy could be recycled to synthesize macromolecules required for gametogenesis [103, 104]. In contrast, the down-regulated processes such as photosynthesis, carbohydrate metabolism occurring during the transition from vegetative cells to gametes are up-regulated after the formation of the zygotes from gametes fusion (**Figure S3B and S3C**).

Taken together, we illustrate the first epitranscriptomic RNA m^6^A profile during sexual reproduction of *Chlamydomonas* and find that t m^6^A exhibits a conservative distribution pattern and is mainly enriched in 3’UTR. More importantly, we found the negative correlation between m^6^A methylation level and gene expression and m^6^A is likely involved in the sexual reproduction through regulating microtubule-based movement. Our study provides new insights into epigenetic regulation in *Chlamydomonas* life cycle, and clues for the further studies of m^6^A modification in the evolution of animal and plant reproduction.

## Materials and methods

### Strains and growth conditions

*Chlamydomonas* strains CC-620 and CC-621 were cultivated on solid TAP medium with photoperiod 12h light/ 12h dark, 22[, 50 μmol photons m^-2^ s^-1^. The cells were resuspended into 60 mL TAP liquid medium with 4∼6×10^7^ cells per mL for further sample collection.

### Sample preparation

Sample preparation was performed as previously described [15] with slight modifications. 15mL *Chlamydomonas* cells was collected as vegetative cell samples at 2 h under light. The remaining cells was resuspended in 45mL TAP-N medium and 15mL of the cells was harvested as gamete cells after culturing in the light for 21h, while the rest of 30mL cells was resuspended in ultrapure water. After being shaken slowly for 30 min under low light, the two strains were mixed in equal amounts, and placed under normal light for 2 h to complete the mating (check the cell mating status under a microscope). The zygotes were transferred to TAP medium containing 3% agar, and half of cells were collected as a 1d zygotes sample after one day in the light, while the remaining half were placed in the dark for 6 days and collected as a 7d zygotes sample. Before collecting the zygotes samples, the lawn on the solid agar surface with vegetative cells was scraped off with a spatula, and the zygotes were collected with a scalpel and resuspended in Tris-EDTA-NaCl (TEN) buffer with 0.2% (v/v) Nonidet P-4 (NP-40) to remove the remaining vegetative cells. After that, the zygotes were collected by centrifugation and resuspended in TEN buffer.

### Total RNA Isolation and mRNA Purification

Total RNA was isolated by Trizol (Invitrogen, 15596026 and 15596018) according to the manufacturer’s protocol. The mRNA of *Chlamydomonas* was purified by the Dynabeads® mRNA Purification Kit (Invitrogen, 61006) according to the manufacturer’s instruction twice to remove the ribosome RNA as much as possible.

### m^6^A Dot-blot assay

m^6^A Dot-blot was performed as described previously [54] with slight modifications. Briefly, the mRNA was serially diluted and loaded as follows: 200ng, 100ng, 50ng, and 25ng. The mRNA was denatured at 70°C for 3 min and transferred to the GE Amersham Hybond-N^+^ membrane (GE, RPN303B) using Bio-Dot Apparatus and a vacuum pump. The mRNA was cross-linked under UV light for 3 min, and the membrane was blocked in PBST with 5% skim milk for 1 h. The membrane with mRNA samples was incubated with diluted anti-m^6^A antibody in PBST with 5% skim milk overnight. The membrane was washed with PBST 3 times for 5 min each time followed the incubation of the diluted goat anti-rabbit secondary antibody in PBST with 5% skim milk for 1 h. After washed with PBST for 3 times, the membrane was incubated with ECL Prime Western Blotting Detection Reagent (GE, RPN2232) for 1 min and was exposed to film. The membrane was stained with methylene blue to check the loading amounts.

### MeRIP-seq

m^6^A MeRIP-seq was performed following the previously reported protocol [105]. The purified mRNA was fragmented to a size about 200 nt using the fragmentation reagent (Life Technologies, AM8740). 30 μL of protein A magnetic beads (Thermo Fisher Scientific, 10002D) was washed twice with 1 mL IP buffer (150 mM NaCl, 10 mM Tris-HCl [pH 7.5], 0.1% NP-40 in nuclease-free H_2_O), then resuspended in 500 μL IP buffer mixed with 5 μg anti-m^6^A antibody (Millipore, ABE572) and incubated at 4 °C with gentle rotation for at least 6 hours. After twice washes with IP buffer (10 mM Tris-HCl [pH 7.4], 150 mM NaCl, 0.1% NP-40 in DEPC-treated), the mixture of antibody and beads was resuspended in 500 μL of the IP reaction mixture containing 500 ng fragmented mRNA, 100 μL of 5× IP buffer, and 5 μL of RNasin Plus RNase Inhibitor (Promega, N2611) and incubated for 2 h at 4 °C. The beads were then washed twice with 1 mL different IPP buffer sequentially, including low-salt IPP buffer (50 mM NaCl, 10 mM Tris-HCl [pH 7.5], 0.1% NP-40 in nuclease-free H_2_O) and high-salt buffer (500 mM NaCl, 10 mM Tris-HCl [pH 7.5], 0.1% NP-40 in nuclease-free H_2_O) for 10 min each time at 4°C. The beads were then eluted with 300 μL IPP buffer with 0.5 mg/mL N^6^-Methyladenosine and RNasin with gentle rotation at room temperature for 1h. The m^6^A-modified RNAs were eluted using 200 μL of RLT buffer supplied in RNeasy Mini Kit (QIAGEN, 74106) for 2 min at room temperature. The supernatant was collected to a new tube and 400 μL of absolute ethanol was added then the mixture was applied to a RNeasy spin column and centrifuged at 13,000 rpm at 4°C for 30 sec. The spin column was washed with 500 μL of RPE buffer supplied in RNeasy Mini Kit, then 500 μL of 80% ethanol, and centrifuged at full speed for 5 min at 4°C to dry the column. The m^6^A-modified RNAs were eluted with 10 μL of nuclease-free H_2_O. For a second round of IP, the eluted RNA was re-incubated with new protein A magnetic beads prepared with new anti-m^6^A antibody, followed by washes, elution and purification as described above. The purified RNAs were used to construct library using the KAPA Standard RNA-Seq Kit according to the manufacturer’s instructions (KAPA, KR1139). The libraries were PCR amplified for 8∼12 cycles and size-selected on the 8% TBE gel. Sequencing was carried out by the Illumina Nova 6000 platform.

### Sequencing data analysis

Pair-end reads with a length of 150 bp were generated by MeRIP-seq and RNA-seq. Cutadapt (version 1.16) [106] software and Trimmomatic (version 0.33) [107] were used to trim off adapter sequences and low quality bases for all of the raw reads. The remaining reads were aligned to the *Chlamydomonas* genome (version 5.6 for assembly; Phytozome version 12 for gene annotation) using Hisat2 (version 2.0.5) [108]. Only uniquely mapped reads with a mapping quality score ≥ 20 were kept for the subsequent analysis for each sample. The number of reads mapped to each gene (Phytozome version 12) was counted using the software featureCounts [109]. For MeRIP-seq, the replicates of each sample were merged (**Figure S1A and S1B**) for m^6^A peak calling using R package exomePeak [110] with the corresponding input samples serving as control. The software BEDTools’ intersectBed (version 2.28.0) [111] was used to annotate each m^6^A peak based on the gene annotation information.

### Statistical analysis of differentially expressed genes and Gene Ontology analysis

Differentially expressed genes among different samples were determined using the R-package edgeR [112]. Transcripts with |log_2_(fold change)|>1 and FDR< 0.01 were considered as significantly differentially expressed genes. Gene Ontology (GO) analysis of specific gene set was performed using agriGO [113] (http://systemsbiology.cau.edu.cn/agriGOv2/). GO terms with *p*-value < 0.05 were statistically significant.

### Motif identification within m^6^A peaks

HOMER (version 4.7) [114] was used to identify the motif enriched by m^6^A peak, and the motif length was limited to 5 nucleotides. The peaks annotated to mRNA were considered as target sequences, and the background sequences were constructed by randomly perturbing these peaks using shuffleBed of BEDTools (version 2.28.0) [111]. Based on the enrichment level, the differential m^6^A peaks were identified with |log2(fold change)| > |log_2_(1.5)|.

### qRT-PCR analysis

RevertAid™ First Strand cDNA Synthesis Kit (Thermo Scientific, K1622) were applied to generate cDNA templates by reverse transcription. The TB Green Premix Ex Taq (TaKaRa, RR420A) was used in the qRT-PCR reaction, and the qRT-PCR was carried out using LightCycler480 (Roche). The *CBLP* gene (Cre06.g278222) was used as the internal control. The calculation of relative mRNA expression was performed as described previously [115].

## Data availability statement

The raw sequence data from RNA-seq and MeRIP-seq have been deposited in the Genome Sequence Archive [116] (Genomics, Proteomics & Bioinformatics 2021) in National Genomics Data Center (Nucleic Acids Res 2021), China National Center for Bioinformation / Beijing Institute of Genomics, Chinese Academy of Sciences (GSA: CRA005106) that are publicly accessible at https://ngdc.cncb.ac.cn/gsa.

## Authors’ contributions

WY conceived this project. YL, HF, GC and YY performed the experiments and analyzed the data. ML, TZ and JW performed bioinformatic analysis. YL, FH, ML, YY and WY wrote the manuscript with input from all authors. All authors read and approved the final manuscript.

## Acknowledgements

This work was supported by National Natural Science Foundation of China (Grant No.91940304), Ministry of Science and Technology of the People’s Republic of China (E0718G3001) and the National Key R&D Program of China (2018YFA0801200).

## Supplementary materials

**Figure S1.**
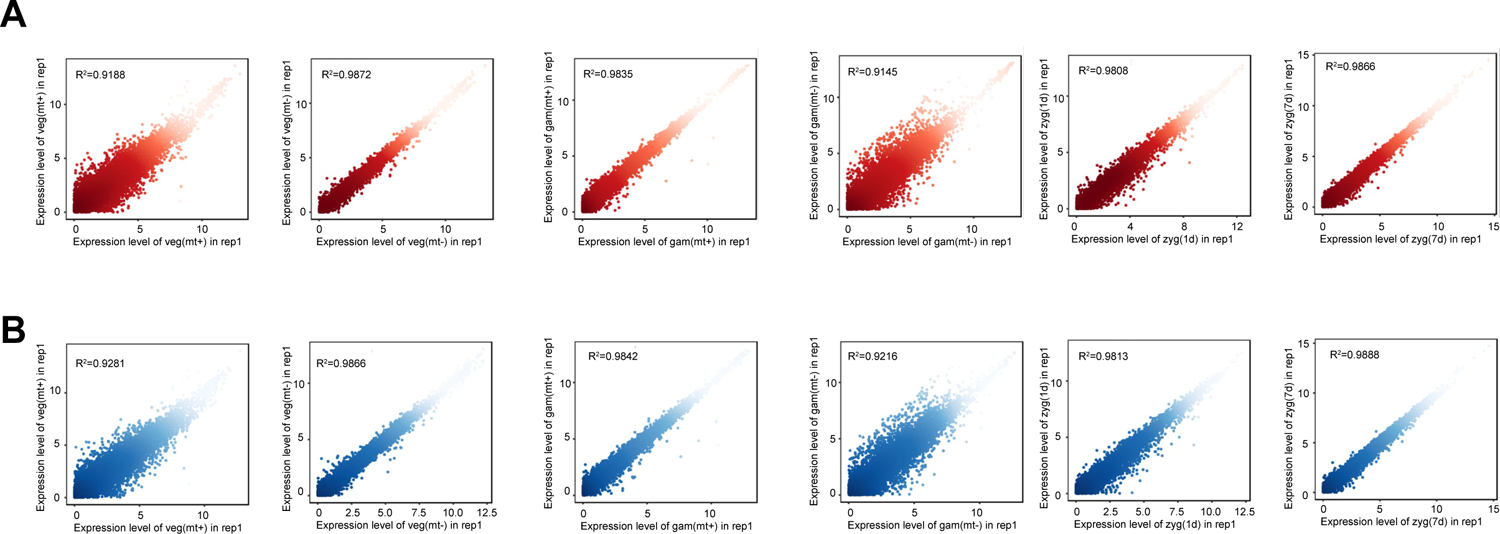
Pearson correlation among the independent biological replicates. **A-B.** Pearson correlation analysis of gene expression among independent biological replicates in RNA-seq (**A**) and MeRIP-seq (**B**), respectively.

**Figure S2.**
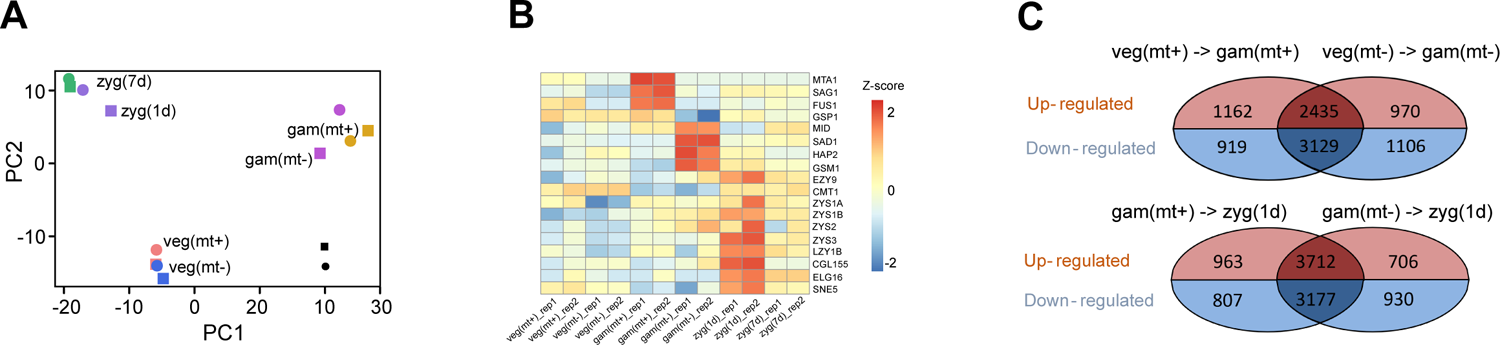
The profiles of gene expression during the life cycle of *Chlamydomonas*. **A.** PCA analysis based on the gene expression exhibiting the high consistency among replicates. **B.** Heatmap showing the stage-specific expression of some of the differentially expressed genes. **C.** Overlap of the up-regulated and down-regulated expressed genes between mating type minus and plus in the processes of vegetative cells to gametes (above) and gametes to zygotes (below).

**Figure S3.**
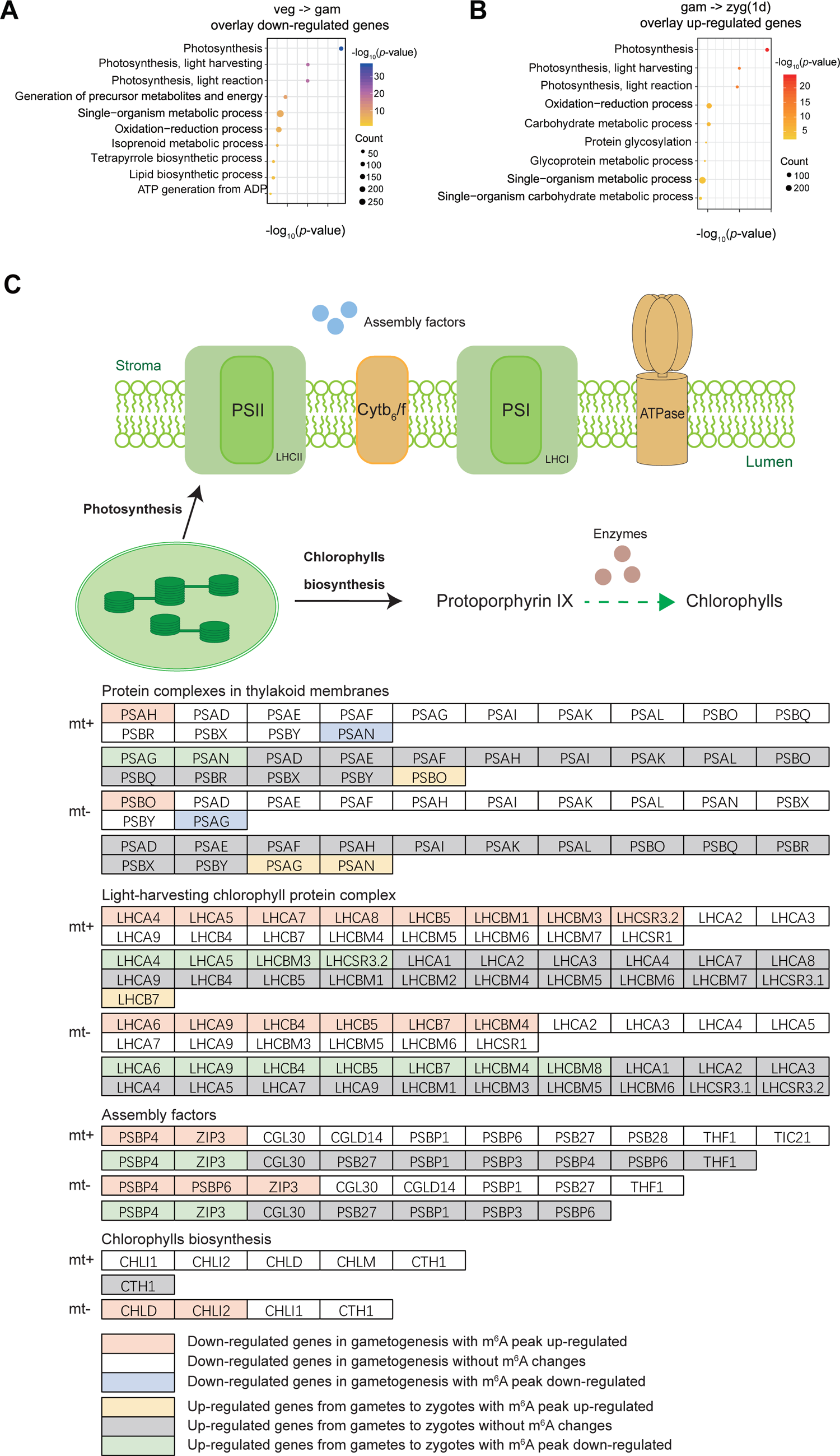
Biological pathways related to the *Chlamydomonas* life cycle. **A.** Gene Ontology (GO) enrichment of down-regulated genes in both gametes with different mating types (mt+ and mt-) during gametogenesis. **B.** GO enrichment of up-regulated genes in 1d zygotes in the process from mating type plus and minus gametes to zygotes. **C.** The m^6^A modification level of photosynthesis associated genes during sexual reproduction.

---

**Table S1** The identified m^6^A peaks at different stages of life cycles

**Table S2** Gene expression RPKM matrix

**Table S3** The differentially expressed genes at different stages

